# Conditional *Pten* knockout in parvalbumin- or somatostatin-positive neurons sufficiently leads to autism-related behavioral phenotypes

**DOI:** 10.1101/2020.04.17.047159

**Authors:** Sangyep Shin, Andrea Santi, Shiyong Huang

## Abstract

Disrupted GABAergic neurons have been extensively described in brain tissues from individuals with autism spectrum disorder (ASD) and animal models for ASD. However, the contribution of these aberrant inhibitory neurons to autism-related behavioral phenotypes is not well understood. We examined ASD-related behaviors in mice with conditional *Pten* knockout in parvalbumin (PV)-expressing or somatostatin (Sst)-expressing neurons, two common subtypes of GABAergic neurons. We found that mice with deletion of *Pten* in either PV-neurons or Sst-neurons displayed social deficits, repetitive behaviors and impaired motor coordination/learning. In addition, mice with one copy of *Pten* deletion in PV-neurons exhibited hyperlocomotion in novel open fields and home cages. We also examined anxiety behaviors and found that mice with *Pten* deletion in Sst-neurons displayed anxiety-like behaviors, while mice with *Pten* deletion in PV-neurons exhibited anxiolytic-like behaviors. These behavioral assessments demonstrate that *Pten* knockout in the subtype of inhibitory neurons sufficiently gives rise to ASD-core behaviors, providing evidence that both PV- and Sst-neurons may play a critical role in ASD symptoms.

## INTRODUCTION

Inhibitory neurons are highly impacted in autism spectrum disorder (ASD) [1–3], a neurodevelopmental condition characterized by symptoms of difficulty in communication, deficits in social interaction, and the presence of restricted/repetitive behaviors [4]. Soma-targeting parvalbumin (PV)-expressing neurons (PV-neurons) and dendritic-targeting somatostatin (Sst)-expressing neurons (Sst-neurons) are two prominent subtypes of inhibitory neurons in the cortex [5]. Studies of postmortem human brain tissues have demonstrated aberrant inhibitory neurons, mainly PV-neurons, in the brains of individuals with autism [6–9]. Disrupted inhibitory neurons have also been extensively reported in animal models for autism [10–12]. However, the contribution of each subtype of inhibitory neurons to autism symptoms is largely unknown. Thus, evaluating the role of PV-neurons and Sst-neurons in autism behaviors is crucial for the development of treatments.

The autism-risk gene, phosphatase and tensin homolog on chromosome ten (*PTEN*), originally identified as a tumor suppressor gene, negatively regulates cell proliferation and growth by downregulating the phosphatidylinositol 3-kinase/AKT/mammalian target of rapamycin (PI3K/AKT/mTOR) pathway [13–16]. *PTEN* germline mutations have been identified in individuals with ASD [17, 18], and may account for up to 12% of ASD cases [19–21]. *Pten* is widely expressed in both glutamatergic and GABAergic neurons during development and adulthood [22, 23]. Since deletion of *Pten* is embryonically lethal, mice with brain-region-specific *Pten* knockout or *Pten* haploinsufficiency have been investigated. These transgenic mice exhibited autism-related behaviors, including deficits in social interaction, repetitive behaviors, hyperlocomotion, and anxiety-like behaviors [24–30]. However, whether conditional knockout of *Pten* in subtypes of inhibitory neurons causes autism-behavioral phenotypes remains elusive.

We used the Cre/loxp recombination system to generate mice with *Pten* knockout in PV-neurons or Sst-neurons, and conducted a battery of behavioral tests to examine autism-related behavioral phenotypes in these mice. We found that conditional knockout *Pten* in PV-neurons or Sst-neurons is sufficient to result in autism-core symptoms, including social deficits and repetitive behaviors. We also showed that *Pten* in PV-neurons and Sst-neurons is crucial for motor coordination and learning. Finally, deletion of *Pten* in PV-neurons induces hyperlocomotion and anxiolytic-like behaviors whereas deletion of *Pten* in Sst-neurons causes anxiety-like behaviors. Collectively, *Pten* mutation in PV-neurons and Sst-neurons results in autism-related behavioral phenotypes.

## RESULTS

### Deletion of Pten in PV-neurons or Sst-neurons leads to social deficits

First, we examined the effect of conditional *Pten* knockout in parvalbumin-expressing neurons (PV-neurons) on social behaviors. We specifically deleted *Pten* in PV-neurons by crossing PV-Cre mice with *Pten*^flox^ mice to generate PV-Cre^+/+^/*Pten*^fl/+^ mice (See Methods for details). Using these mice as breeders, we obtained offspring littermates with three genotypes, PV-Cre^+/+^/*Pten*^+/+^ (PV-Pten-WT), PV-Cre^+/+^/*Pten*^fl/+^ (PV-Pten-Het), and PV-Cre^+/+^/*Pten*^fl/fl^ (PV-Pten-KO). These three types of mice all have homozygous Cre but different copies of *Pten* in PV-neurons. Using immunohistochemistry and western blot, we confirmed that *Pten* was specifically reduced and deleted in PV-positive neurons of PV-Pten-Het and PV-Pten-KO mice, respectively (Figure S1). Sociability and social novelty preference in these mice were assessed using modified three-chamber social tests (Figure 1A). Firstly, we confirmed that there was no pre-existing side preference in the PV-Pten mice (Figure S3A). As shown in figure 1B, single or two copies of *Pten* deletion in PV-neurons did not change the social approach behaviors. All three genotypes of PV-Pten mice spent more time with social partners than with objects in the sociability test. The preference index was not different among the three groups of PV-Pten mice (Figure 1B). In contrast, knockdown or knockout of *Pten* in PV-neurons impaired the social novelty preference. In the social novelty test, only PV-Pten-WT mice spent more time interacting with novel social partners (Figure 1C). The preference index for social novelty decreased in PV-Pten-Het and PV-Pten-KO mice compared to that in PV-Pten-WT mice (Figure 1C).

**Figure 1.**
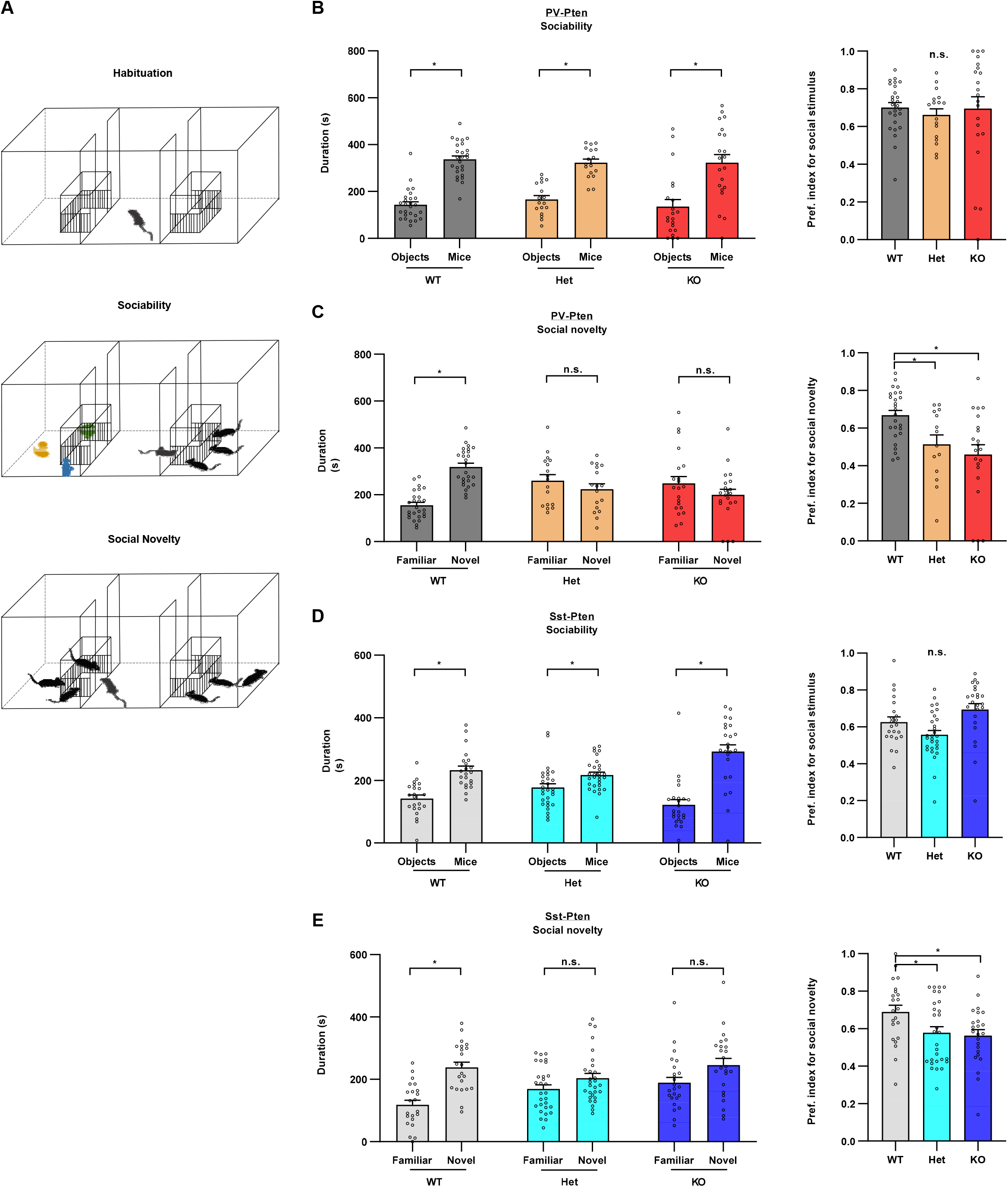
PV-Pten-KO and Sst-Pten-KO mice displayed social deficits in the modified three-chamber social test. **(A)** Schematic diagrams of the modified three-chamber social test. (**B**) All three genotypes of PV-Pten mice showed significant social preference between objects and mice (social partners) (WT: n = 26, W = 325, *p* < 0.0001, Wilcoxon test; Het: n = 17, *t*_(16)_ = 5.065, *p* = 0.0001, paired *t*-test; KO: n = 21, W = 151, *p* = 0.0071, Wilcoxon test). There is no significant difference in preference index for social stimulus among these three genotypes of PV-Pten mice (*F*_(2,61)_ = 0.213, *p* = 0.809, one-way ANOVA). (**C**) Only PV-Pten-WT mice displayed significant preference between familiar and novel mice in the social novelty test (WT: n = 26, *t*_(25)_ = 6.142, *p* < 0.0001; Het: n = 17, *t*_(16)_ = 0.7481, *p* = 0.4652; KO: n = 21, *t*_(20)_ = 0.9664, *p* = 0.3454; Paired *t*-test). The preference index for social novelty significantly reduced in PV-Pten-Het and PV-Pten-KO mice compared with PV-Pten-WT mice (*F*_(2,61)_ = 7.402, *p* = 0.0011, one-way ANOVA; Het vs. WT: *p* = 0.0427, KO vs. WT: *p* = 0.0011, Tukey’s post hoc test). (**D**) All three genotypes of Sst-Pten mice showed significant social preference between objects and social partners (WT: n = 22, *t*_(21)_ = 4.570, *p* = 0.0002, paired *t*-test; Het: n = 29, W = 221.0, *p* = 0.0157, Wilcoxon test; KO: n = 24, W = 254.0, *p* < 0.0001; Wilcoxon test). The preference index for social stimulus reduced slightly in Sst-Pten-Het mice and increased in Sst-Pten-KO mice, but none of these changes reached statistical significance in post hoc comparisons (*p* = 0.0008, Kruskal-Wallis test; Het vs. WT: *p* = 0.2287, KO vs. WT: *p* = 0.0855, Dunn’s post hoc test). (**E**) Sst-Pten-Het and Sst-Pten-KO mice displayed deficits in the social novelty test (WT: n = 22, *t*_(21)_ = 5.269, *p* < 0.0001, paired *t*-test; Het: n = 29, W = 99, *p* = 0.2941, Wilcoxon test; KO: n = 24, *t*_(23)_ = 1.775, *p* = 0.0891, paired *t*-test). The preference index for social novelty significantly reduced in Sst-Pten-Het and Sst-Pten-KO mice (*p* = 0.0311, Kruskal-Wallis test; Het vs. WT: *p* = 0.0442, KO vs. WT: *p* = 0.0384, Dunn’s post hoc test). The preference index was calculated as described in the methods. Circles represent data from individual animals, and bar graphs indicate mean + SEM. *: significant; n.s.: not significant.

Next, we evaluated the effect of conditional *Pten* knockout in somatostatin-expressing neurons (Sst-neurons) on social behaviors. Using the same breeding strategy for PV-Pten mice, we obtained Sst-Pten littermates with three genotypes, Sst-Cre^+/+^/*Pten*^+/+^ (Sst-Pten-WT), Sst-Cre^+/+^/*Pten*^fl/+^ (Sst-Pten-Het), and Sst-Cre^+/+^/*Pten*^fl/fl^ (Sst-Pten-KO). We also verified that *Pten* was specifically reduced and deleted in Sst-positive neurons of Sst-Pten-Het and Sst-Pten-KO mice, respectively (Figure S2), and there was no pre-existing side preference in the Sst-Pten mice (Figure S3B). We found that all three genotypes of Sst-Pten mice spent more time in the interaction areas for social partners in the sociability test (Figure 1D). This result was also confirmed by the preference index for sociability which was slightly reduced in Sst-Pten-Het mice while increased in Sst-Pten-KO mice, but these changes were not significant in post hoc comparisons (Figure 1D). In the social novelty test, only Sst-Pten-WT but not -Het or -KO mice spent more time in the interaction zones for novel social partners (Figure 1E). Consistently, the preference indexes for social novelty was reduced in both Sst-Pten-Het and -KO mice (Figure 1E). Thus, Sst-Pten-Het and -KO mice displayed normal sociability but exhibited impairment in social novelty behavior. Collectively, elimination of *Pten* in either PV-neurons or Sst-neurons resulted in reduced social behaviors, particularly social novelty preference.

### Deficits of locomotion and motor coordination and increased repetitive behaviors in PV-Pten and Sst-Pten mice

We then examined the contribution of *Pten* mutations in PV- and Sst-neurons to deficits in locomotor behaviors. 30-min open field tests were performed in PV-Pten mice (Figure 2A). PV-Pten-Het but not -KO mice traveled significantly longer distances compared to PV-Pten-WT (Figure 2B). When examined in 5-min intervals, the distance moved in PV-Pten-WT and -Het mice declined slightly over the 30-min testing period, and PV-Pten-Het consistently traveled longer distances in each interval. PV-Pten-KO mice maintained the same rate of activity over the 30-min testing time (p = 0.903, Pearson correlation test), but no statistical difference compared to PV-Pten-WT mice (Figure 2C). We examined the locomotor activity in home cages as well. Consistent to the result in the open-field tests, PV-Pten-Het mice but not PV-Pten-KO mice exhibited increased locomotor activity (Figure 2D and 2E). Thus, PV-Pten-Het mice exhibited elevated locomotor activity both in the novel open field and home cages. In Sst-Pten mice, the locomotor activity was not different among genotypes (Figure 2F), and was confirmed by measurements across 5-min intervals (Figure 2G). Sst-Pten mice also performed normally in the home-cage activity test (Figure 2H–2I).

**Figure 2.**
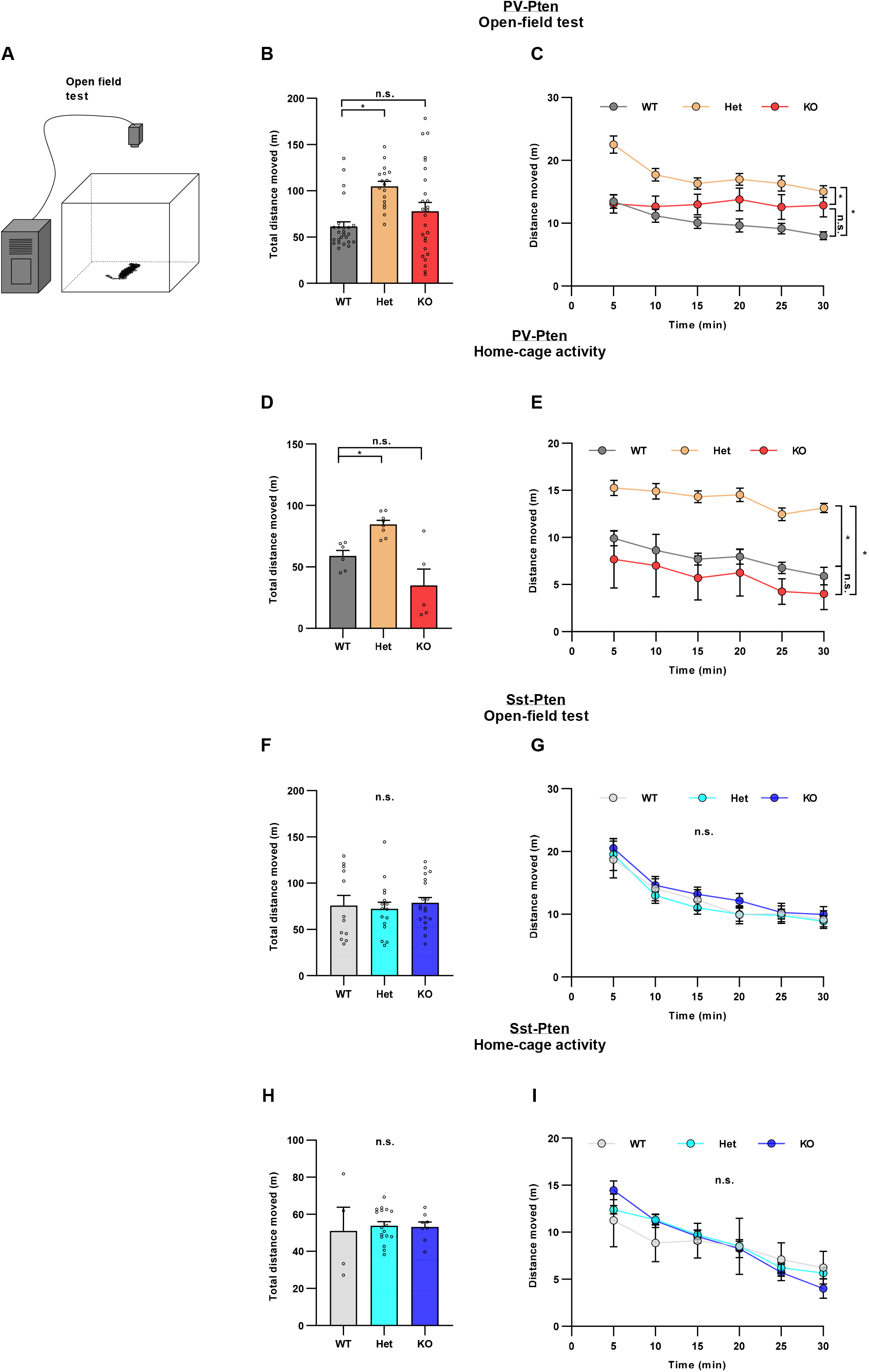
PV-Pten-Het exhibited hyper-locomotion in the open field test. (**A**) The schematic diagram of the open field test. (**B**) PV-Pten-Het mice traveled a longer distance than PV-Pten-WT mice during the 30-min test (WT: n = 25, Het: n = 17, KO; n = 26; *F*_(2,65)_ = 7.244, *p* = 0.001, one-way ANOVA; Het vs. WT: *p* < 0.001, KO vs. WT: *p* = 0.236, Tukey’s post hoc test). (**C**) When the total distance of PV-Pten mice was analyzed in 5-min time bin, PV-Pten-Het mice displayed hyperlocomotion (*p* = 0.0012, genotype effect, two-way rmANOVA; Het vs. WT: *p* < 0.001, KO vs. WT: *p* = 0.236, Tukey’s post hoc test). (**D**) PV-Pten-Het mice exhibited increased travel distance in 30-min home-cage activity (WT: n = 6, Het: n = 8, KO; n = 5; *F*_(2,16)_ = 12.93, *p* = 0.0005, one-way ANOVA; Het vs. WT: *p* = 0.036; KO vs. WT, *p* = 0.087, Tukey’s post hoc test). (**E**) Home-cage activity was analyzed in a 5-min time bin. PV-Pten-Het mice displayed hyperlocomotion in home cages (*p* = 0.0001, genotype effect, two-way rmANOVA; Het vs. WT: *p* = 0.0017, KO vs. WT: *p* = 0.4685, Tukey’s post hoc test). (**F-G**) There is no significant difference in total distance moved (D; WT: n = 12, Het: n = 16, KO: n = 19; *F*_(2,44)_ = 0.313, *p* = 0.733, one-way ANOVA) or distance moved in 5-min time bin (E) in Sst-Pten mice (*p* = 0.7002, genotype effect, two-way rmANOVA). **(H-I)** In home-cage activity test, there was no difference in the total distance traveled (H; WT: n = 4, Het: n = 17, KO: n = 8; *F*_(2,26)_ = 0.088, *p* = 0.9164, one-way ANOVA) or the distance moved in 5-min time bin (I; *p* = 0.9164, genotype effect, two-way rmANOVA) among the three genotypes of Sst-Pten mice. In B, D, F and H, circles represent data from individual animals, and bar graphs indicate mean + SEM. In C, E, G and I, data are represented as mean ± SEM. *: significant; n.s.: not significant.

We next examined self-grooming during the open field test for repetitive behaviors. PV-Pten-KO mice exhibited an increase of grooming bouts (Figure 3A) and spent more time in self-grooming in the open field (Figure 3B). Similarly, the number of grooming bouts and total time of self-grooming were increased in Sst-Pten-KO mice (Figure 3C and 3D). These data suggest that deletion of *Pten* in PV-neurons or Sst-neurons induces repetitive behaviors, a core-domain of autism behaviors.

**Figure 3.**
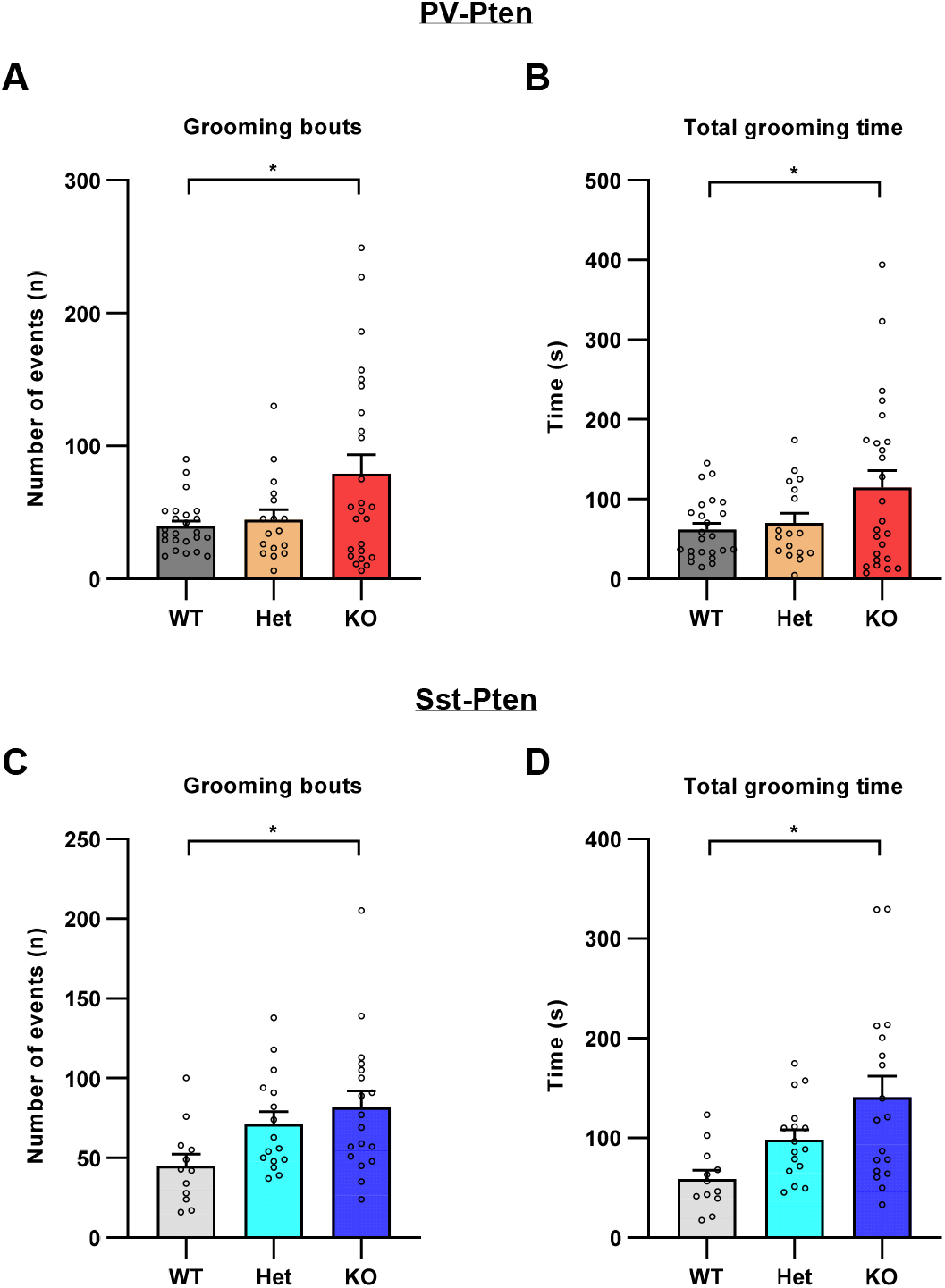
Increased self-grooming in PV-Pten-KO and Sst-Pten mice. (**A**) The number of self-grooming bouts increased in PV-Pten-KO mice (WT: n = 24, Het: n = 17, KO: n = 25; *F*_(2,63)_ = 4.758, *p* = 0.012, one-way ANOVA; KO vs. WT: *p* = 0.015, Tukey’s post hoc test). (**B**) The total time of self-grooming increased in PV-Pten-KO mice (WT: n = 24, Het: n = 17, KO: n = 25; *F*_(2,63)_ =3.635, *p* = 0.032, one-way ANOVA; KO vs. WT: *p* = 0.036, Tukey’s post hoc test). (**C**) The number of self-grooming bouts increased in Sst-Pten-KO mice (WT: n = 12, Het: n = 16, KO: n = 18; *F*_(2,43)_ = 4.025, *p* = 0.0250, one-way ANOVA; KO vs. WT: *p* = 0.0199, Tukey’s post hoc test). (**D**) The total time of self-grooming increased in Sst-Pten-KO mice (WT: n = 12, Het: n = 16, KO: n = 18; *F*_(2,43)_ = 6.249, *p* = 0.004, one-way ANOVA; KO vs. WT: *p* = 0.003, Tukey’s post hoc test). Circles represent data individual animals, and bar graphs indicate mean + SEM. *: statistically significant.

We further evaluated the motor coordination/learning in PV-Pten mice using the rotarod test (Figure 4A). Mice were trained on an accelerating rotarod for 6 trials over two days (3 trials per day). The latency to fall was significantly reduced in PV-Pten-Het and PV-Pten-KO mice over the 6-trial training (Figure 4B). Particularly, motor coordination was severely impaired in PV-Pten-KO mice. They were barely able to stay on the stationary rotarod (Figure 4B). We next analyzed the learning index (ratio of latency to fall between the 6^th^ trial and the 1^st^ trial) and found that knockout of *Pten* in PV-neurons impaired motor learning in the rotarod test (Figure 4C). Similarly, knockout of *Pten* in Sst-neurons impaired rotarod performance and motor learning as well (Figure 4D and 4E). Collectively, knockdown or knockout of *Pten* in PV-neurons or Sst-neurons impairs motor coordination and learning.

**Figure 4.**
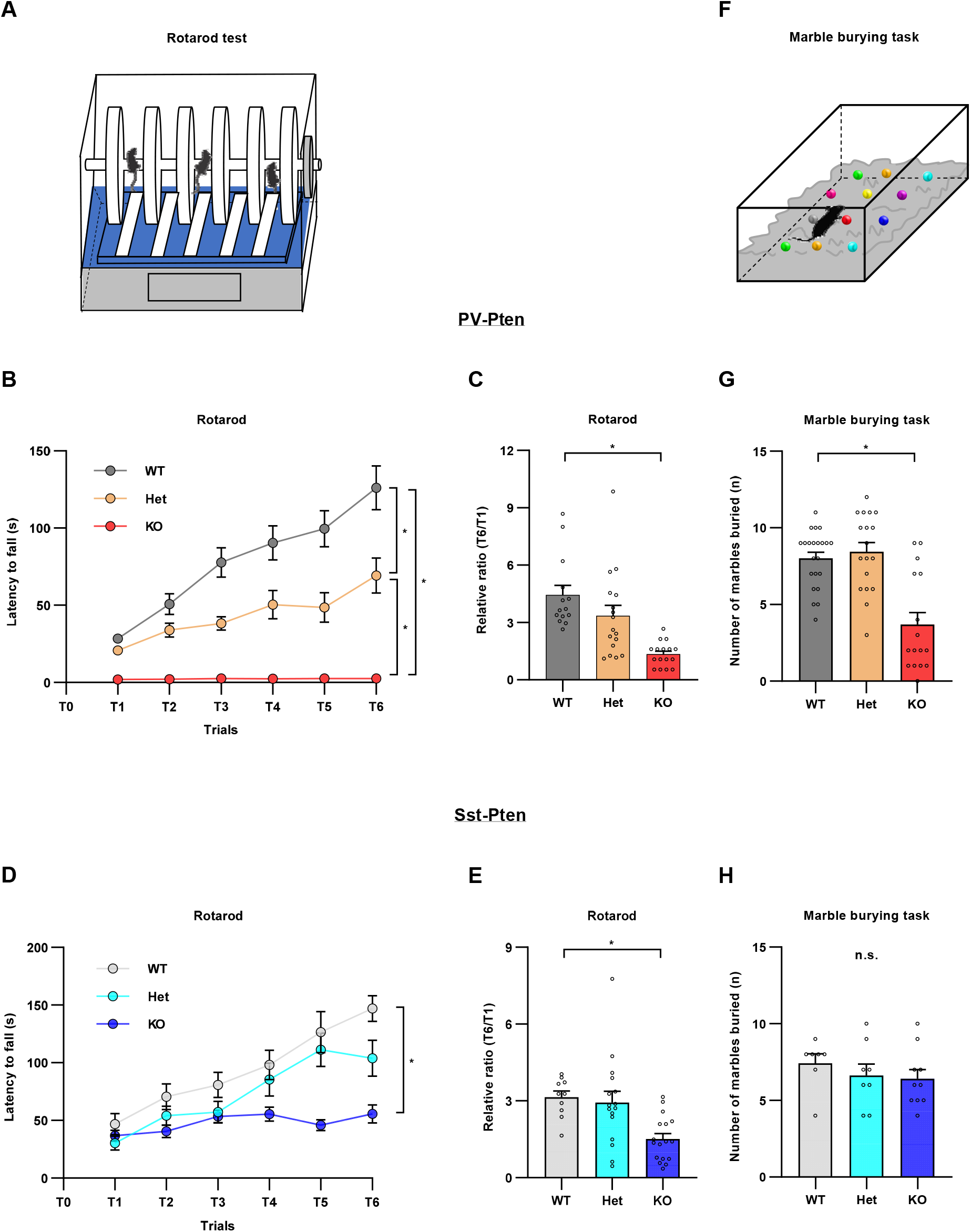
Impaired motor function in PV-Pten-KO and Sst-Pten-KO mice. (**A**) The schematic diagram of the rotarod test.(**B**) PV-Pten-Het and PV-Pten-KO mice showed a shorter latency to fall off the rotarod than PV-Pten-WT over the 6-trials training (WT: n = 14, Het: n = 17, KO: n = 14; *p* < 0.0001, genotype effect, two-way rmANOVA; Het vs. WT: *p* < 0.0001, KO vs. WT: *p* < 0.0001, Het vs. KO: *p* < 0.0001, Tukey’s post hoc test). (**C**) The learning index (T6/T1) was significantly reduced in PV-Pten-KO mice (*F*_(2,44)_ = 12.435, *p* < 0.001, one-way ANOVA; KO vs. WT: *p* < 0.001, Tukey’s post hoc test). (**D**) Sst-Pten-KO mice showed a shorter latency to fall off the rotarod than Sst-Pten-WT mice over the 6-trials training (WT: n = 10, Het: n = 16, KO: n = 17; *p* = 0.0005, genotype effect, two-way rmANOVA; KO vs. WT: *p* = 0.0004, Tukey’s post hoc test). (**E**) The learning index (T6/T1) was significantly reduced in Sst-Pten-KO mice (*F*_(2,40)_ = 7.372, *p* = 0.0019, one-way ANOVA; KO vs. WT: *p* = 0.0065, Tukey’s post hoc test). (**F**) The schematic diagram of the marble burying test. (**G**) PV-Pten-KO mice buried significantly fewer marbles compared to PV-Pten-WT mice (WT: n = 22, Het: n = 18, KO: n = 16; *p* < 0.001, Kruskal-Wallis test; KO vs. WT, *p* = 0.001, Dunn’s post hoc test). (**H**) There is no difference in the number of marbles buried among the three genotypes of Sst-Pten mice (WT: n = 7, Het: n = 8, KO: n = 10; *p* = 0.406, Kruskal-Wallis test). In C, E, G and H, circles represent data from individual animals, and bar graphs indicate mean + SEM. In B and D, data are represented as mean ± SEM. *: significant; n.s.: not significant.

We then conducted marble burying tests in PV-Pten and Sst-Pten mice (Figure 4F). PV-Pten-KO mice buried significantly fewer marbles than PV-Pten-WT mice did (Figure 4G), but there was no difference in the marble burying task among the three genotypes of Sst-Pten mice (Figure 4H). Since the burying of marbles entails physical motor activity, reduced marble burying activity in PV-Pten-KO mice may be mainly resulted from impaired motor coordination in this type of mice (Figure 4B).

### Anxiolytic-like behavior in PV-Pten mice and anxiety-like behavior in Sst-Pten mice

Finally, we examined anxiety-like behaviors in mice with *Pten* deletion in inhibitory neurons using elevated plus maze tests (Figure 5A) and open field tests (Figure 5B). In the elevated plus maze, PV-Pten-KO mice spent increased percentage of time in the open arms and decreased percentage of time in the closed arms, compared to PV-Pten-WT and PV-Pten-Het mice (Figure 5C and 5D). Consistently, in the open field test, PV-Pten-KO mice spent increased time in the center area (Figure 5E), and decreased time in the corner area of the open field compared to PV-Pten-WT mice (Figure 5F). In contrast, Sst-Pten-KO mice showed a reduced percentage of time in the open arms and an increased percentage of time in the closed arms (Figure 5G and 5H). We did not observe any difference of time spent in the center or corner areas among the three genotypes of Sst-Pten mice (Figure 5I and 5J). Thus, we concluded that *Pten* knockout in PV-neurons induces anxiolytic-like behavior while *Pten* knockout in Sst-neurons evokes anxiety-like behavior.

**Figure 5.**
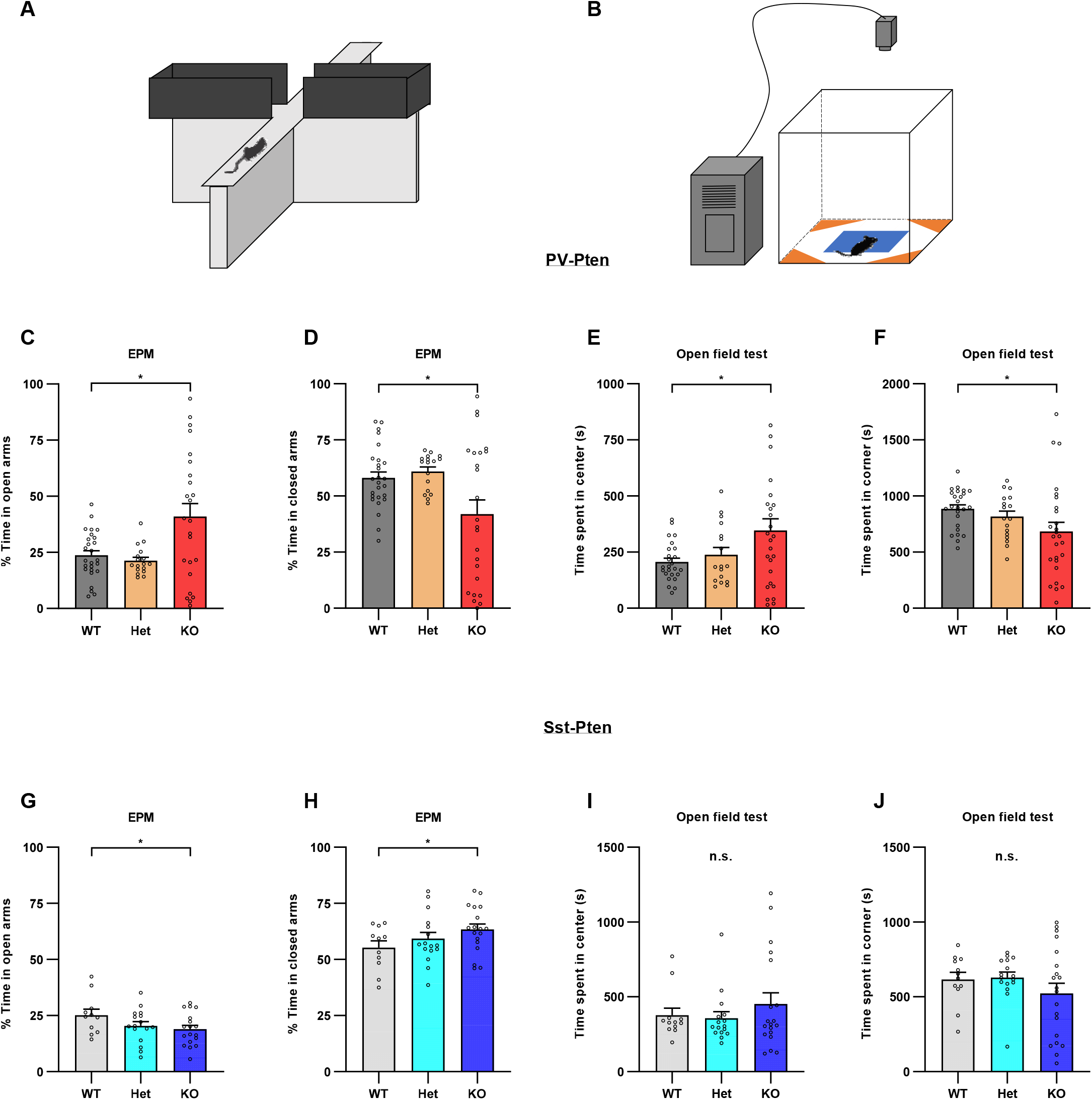
PV-Pten-KO mice showed anxiolytic behaviors while Sst-Pten-KO mice displayed anxiety-like behaviors, compared to their WT mice. (**A**) The schematic diagram of the elevated plus-maze test (EPM). (**B**) The schematic diagram of the center (blue) and corner area (orange) in the open field test. (**C**) PV-Pten-KO mice spend more percentage of time in the open arms compared to PV-Pten-WT and PV-Pten-Het mice (WT: n = 26, Het: n = 17, KO: n = 24; *F*_(2,64)_ = 7.373, *p* = 0.001, one-way ANOVA; KO vs. WT: *p* = 0.005, Tukey’s post hoc test). (**D**) PV-Pten-KO mice spend less percentage of time in the closed arms compared to PV-Pten-WT and PV-Pten-Het mice (*F*_(2,64)_ = 5.310, *p* = 0.007, one-way ANOVA; KO vs. WT: *p* = 0.023, Tukey’s post hoc test). (**E-F**) In the open field test, PV-Pten-KO mice spent more time in the center area (E) and less time in the corner area (F), compared to PV-Pten-WT mice (WT: n = 25, Het: n = 17, KO: n = 26; Center: *p* = 0.024, one-way ANOVA; KO vs. WT: *p* = 0.023, Tukey’s post hoc test; Corner: *p* = 0.016, Kruskal-Wallis test; KO vs. WT: *p* = 0.009, Dunn’s post hoc test). (**G**) Sst-Pten-KO mice spend less percentage of time in the open arms compared to Sst-Pten-WT mice (WT: n = 11, Het: n = 16, KO: n = 17; *F*_(2,34)_ = 3.984, *p* = 0.028, one-way ANOVA; KO vs. WT: *p* = 0.026, Tukey’s post hoc test). (**H**) Sst-Pten-KO spend more percentage of time in the closed arms compared to Sst-Pten-WT mice (*F*_(2,34)_ = 3.596, *p* = 0.038, one-way ANOVA; KO vs. WT: *p* = 0.031, Tukey’s post hoc test). (**I-J**) In the open field test, there is no difference in time spent in the center (I) or corner area (J) among the three genotypes of Sst-Pten mice (WT: n = 12, Het: n = 16, KO: n = 19; Center: *p* = 0.851, Kruskal-Wallis test; Corner: *p* = 0.502, Kruskal-Wallis test). Circles represent data from individual animals, and bar graphs indicate mean + SEM. *: significant; n.s.: not significant.

## DISCUSSION

Our study revealed that conditional deletion of *Pten* in either PV-neurons or Sst-neurons results in autism-like core symptoms, including reduced social novelty preference and presence of repetitive behaviors (self-grooming). In addition, conditional *Pten* knockout in PV-neurons causes differences in behaviors that are characteristic symptoms often observed in autism, such as hyperactivity and impairment in motor coordination/learning. Similarly, conditional *Pten* knockout in Sst-neurons gives rise to disrupted motor coordination/learning. Interestingly, PV-Pten-KO mice displayed anxiolytic-like behaviors, whereas Sst-Pten-KO mice showed anxiety-like behaviors. These findings provide the behavioral evidence that *Pten* mutation in PV-neurons or Sst-neurons sufficiently induces ASD-related behaviors.

The mice used in this study all had homozygous Cre and different copies of *Pten* in PV-neurons or Sst-neurons. Reduced somatostatin expression in homozygous Sst-Cre mice has been previously reported [31], which might potentially impact behavioral performance. We compared behaviors in PV-Pten-WT mice and Sst-Pten-WT mice with those in Pten^fl/fl^ mice, and found that PV-Pten-WT and Sst-Pten-WT mice did not exhibit any difference in most behavioral tests, except that Sst-Pten-WT mice displayed increased time spent in the center of the open field test (Figure S4). Since Cre expression is just the background for all genotypes of mice used here, our conclusion in this study should not be affected by the change of baseline behavioral performance in Sst-Pten-WT mice. We were also wondering whether the different ages of mice performed similarly in the behavioral tests. By performing correlation tests, we found that there was no relationship between the age and behavioral performance in all behavioral tests (Figure S5 and S6). Additionally, the prevalence of autism is different in sex [32] and our previous study in animal models also showed the sex difference in behaviors of *Slit3*-KO mice [33]. However, we did not found any sex effect in the behavioral phenotypes of PV-Pten and Sst-Pten mice (Figure S7 and S8).

Mice with deletion of several other genes (*Shank2*, *mGluR5* and *Lom4*) in PV-neurons exhibit autism-related behaviors of varying severity [34–36], supporting that the function of PV-neurons is an important determinant of autism behaviors. Autism-relevant behavioral phenotypes are also reported in mice with conditional knockout of *MeCP2* (Rett syndrome) or *Scn1a* (Dravet syndrome) in PV-neurons but not Sst-neurons [37, 38]. In our results, conditional knockout of *Pten* in either PV- or Sst-neurons resulted in autism-like core behaviors, further suggesting that *Pten* plays an important role in the function of these two subtypes of neurons.

Three-chamber social tests are commonly used to examine sociability and social novelty preference in mice [39]. The sociability test is to assess the tendency of initiating social contact and remaining proximal to an unfamiliar conspecific, whereas the social novelty test is to evaluate the tendency of approaching novel mice, the discrimination between familiar and novel mice, and the recognition/memorization of the mouse that has interacted before [40]. PV-Pten-KO and Sst-Pten-KO mice exhibited deficits mainly in social novelty preference. Multiple brain regions could be involved in the social deficits observed in PV-Pten-KO and Sst-Pten-KO mice [41]. The prefrontal cortex (PFC) is one of such brain structures that are well established for social behaviors [42]. PV-neurons and Sst-neurons in medial prefrontal cortex (mPFC) specifically respond to social stimuli [43, 44]. Social-evoked responses in PV-neurons are diminished in *Cntnap2*-KO mice which display deficits in sociability [43]. Consistently, optogenetic activation of PV-neurons in mPFC rescues deficits of sociability and social novelty preference in autism mouse models [43, 45]. However, evidence supporting a relationship between PV-neurons in mPFC and social deficits is not unequivocal [46].

The hippocampus is another brain region which is involved in social interaction, particular social memory, a process that is critical for social novelty preference [47, 48]. PV-neurons in the ventral hippocampus (vHipp) exhibited increased activity to novel social partners compared to familiar ones, and inactivation of PV-neurons but not Sst-neurons in vHipp disrupts social novelty [49]. Conversely, another study reported that knockdown Sst but not PV in vHipp disrupted social interaction [50]. Our data showed that *Pten* deletion in both PV- and Sst-neurons are important for social novelty, indicating that multiple brain regions might contribute to social novelty preference.

The cerebellum was recently implicated in social behaviors [51, 52]. PV and Sst are expressed in Purkinje cells (PC) of the cerebellum [53–55]. Abnormal PC function has been linked to social deficits in mice with conditional deletion of autism-risk genes in PCs [56–59]. However, conditional *Pten* knockout in Purkinje cells results in autism-related behaviors, such as less self-grooming and reduced sociability [28], different phenotypes to those exhibited in PV-Pten-KO mice (increased self-grooming and reduced social novelty preference). This discrepancy may be explained by the fact that Cre-recombinase activity in PV-Cre mice is not only present in Purkinje cells but also in cerebellar interneurons [53, 60], indicating that cerebellar interneurons may contribute to behavioral phenotypes in PV-Pten-KO mice as well. In addition, other brain regions, such as the amygdala [61, 62], ventral tegmental area (VTA)-to nucleus accumbens (NAc) projection [63], and the striatum [64, 65] all have been reported to be involved in social behaviors.

Mutation of *Pten* in PV-neurons may contribute to hyperactivity, a common comorbidity of autism [66]. However, only PV-Pten-Het mice exhibited hyperactivity in the open field test. Both PV-Pten-KO mice and Sst-Pten-KO mice displayed impaired motor coordination/learning in accelerating rotarod tasks and increased repetitive behaviors (self-grooming) in open field tests. Especially, PV-Pten-KO mice were barely able to stay on the rotarod. This severe deficit of motor coordination in PV-Pten-KO mice may mask the hyperactivity resulting from the *Pten* knockout in PV-neurons. The cerebellum and striatum, two brain structures critical for motor coordination/learning and repetitive behaviors [67–69], may account for these behavioral phenotypes in PV-Pten and Sst-Pten mice. Additionally, *Pten* knockout in Purkinje cells results in impaired motor coordination/learning [28], and inhibition of PCs in cerebellar right crus I results in repetitive behaviors [59]. Moreover, *Pten* regulates striatal dopamine signaling [70, 71]. All these lines of evidence indicate that motor deficits and repetitive behaviors in PV-Pten and Sst-Pten mice may result from the functional change of PV- and Sst-neurons in the cerebellum and/or striatum; however, further investigations are needed.

Our results showed that PV-Pten-KO mice exhibited anxiolytic-like behaviors while Sst-Pten-KO mice displayed anxiety-like behaviors. Activation of PV-positive neurons in the dentate gyrus results in anxiolytic behaviors [72]. Interestingly, PV activity in the nucleus accumbens shell (sNAc) negatively correlates with the time of open-arm exploration in EPM test [73]. The mechanisms underlying the differential effects of *Pten* knockout in these two subtypes of neurons on anxiety-like behaviors are not clear.

At the synaptic level, conditional knockout *Pten* from GABAergic cortical interneurons increases the synaptic input onto interneurons [74] and the inhibitory output onto glutamatergic neurons [22, 75]. Conversely, a single copy of *Pten* deletion from PV-neurons impairs the formation of perisomatic inhibition [76]. Though the underlying synaptic mechanisms in PV-Pten-KO and Sst-Pten-KO mice need further investigation, our results provide behavioral evidence that deletion of *Pten* in either subtype of inhibitory neurons elicits autism-related phenotypes, suggesting that a potential benefit from therapeutic strategies targeting either PV-neurons, Sst-neurons, or both subtypes, depending on specific alterations in behavioral phenotype.

## MATERIAL AND METHODS

### Animals

*Pten*^fl/+^, Sst-Cre^+/-^ and PV-Cre^+/-^ mice were purchased from Jaxson Laboratory (Stock #: 006440, 013044 and 008069, respectively). All these strains were maintained on C57BL6/J background. We crossed Pten^fl/fl^ mice with Sst-Cre^+/+^ mice to generate Sst-Cre^+/-^/Pten^fl/+^ mice which were then backcrossed with Sst-Cre^+/+^ mice to generate Sst-Cre^+/+^/*Pten*^fl/+^ breeders. By crossing the breeders we obtained three types of experimental mice, Sst-Cre^+/+^/*Pten*^+/+^ (Sst-Pten-WT), Sst-Cre^+/+^/*Pten*^fl/+^ (Sst-Pten-Het) and Sst-Cre^+/+^/*Pten*^fl/fl^ (Sst-Pten-KO) mice. They were all homozygous for Cre but expressed two, one or no copy of *Pten* in Sst neurons. Using the same breeding strategy, we generated PV-Cre^+/+^/Pten^fl/+^ breeders and their offspring, PV-Cre^+/+^/*Pten*^+/+^ (PV-Pten-WT), PV-Cre^+/+^/*Pten*^fl/+^ (PV-Pten-Het) and PV-Cre^+/+^/*Pten*^fl/fl^ (PV-Pten-KO) mice. Both male and female littermates at the age of 4 - 8 weeks were used for behavioral assessments. There was no significant correlation between the behavioral performance and ages in PV-Pten or Sst-Pten WT mice (Figure S5 and S6). No significant sex effects were observed (Figure S7 and S8). Age and sex-matched C57BL/6 mice (purchased from Veterinary Resources at the University of Maryland School of Medicine) were used as social partners in modified three-chamber social tests. All mice were housed 2 - 5 per cage in ventilated racks in a temperature- and humidity-controlled animal room on a 12 h light/dark cycle with lights on from 07:00 to 19:00 and cared by the AAALAC accredited program of the University of Maryland School of Medicine. Autoclaved rodent chow and water were available *ad libitum*. The experimental protocol was approved by the Institutional Animal Care and Use Committees at the Hussman Institute for Autism and the University of Maryland School of Medicine.

### Immunohistochemistry

Animals were anesthetized with isoflurane and perfused transcardially with 0.9% saline solution. Brains were fixed in 4% paraformaldehyde. Double immunohistochemistry with either anti-parvalbumin made in guinea pig (1:3000 dilution; Swant; Catalog # GP72) or anti-somatostatin made in rat (1:200 dilution; MilliporeSigma Cat.# MAB354), and anti-PTEN made in rabbit (1:250 dilution; Cell Signaling; Catalog # 9559T) were performed as described [77] in coronal sections of 40 μm obtained with a vibratome (Leica VT 1000S). Appropriate Alexa fluor conjugated antibodies (Invitrogen) were used to detect primary antibodies. A negative control with normal goat serum in place of primary antibodies was performed on adjacent sections. Images were obtained with LSM 780 (Zeiss) confocal microscope. Imaging conditions were maintained over the different genotypes within each strain. Cells were counted in cortical layers III-IV (PV^+^ cells) or V-VI (Sst^+^) from matching sections (Bregma -1.7 mm) of three mice per genotype.

PTEN expression levels were quantified by measuring the fluorescence intensity in parvalbumin or somatostatin positive cells with Fiji image analysis package [78]. After background subtraction, fluorescence intensities were normalized by the average PTEN fluorescence intensity in WT mice.

### Western Blot

Cerebellum (PV-Pten mice) or cortex (Sst-Pten mice) samples were homogenized in lysis buffer (PH 7.4; 10 mM Tris HCl, 150 mM NaCl, 1 mM EDTA, 1% Triton X-100, 0.5% NP-40, 1 mM sodium orthovanadate, 1 mM PMSF) with protease inhibitor (Halt protease inhibitor, Pierce). 50 μg of total protein was separated in 4-20% mini-Protean TGX Stain-Free precast gels (BioRad) and transferred to nitrocellulose membranes. Antibodies used included rabbit anti-PTEN, mouse anti β-Actin (Cell Signaling), and rabbit (Cell Signaling) and mouse (Novex) HRP-conjugated secondaries.

### Behavioral Assessments

Following procedures established in our previous work [33], a series of behavioral assessments were conducted. Due to poor reproductive performance of mutant mice, multiple behavioral assessments were performed on the same groups of mice. Each mouse was tested for only one behavioral assessment per day. The behavioral tests were performed in the order of potential stress elicited from the tasks during the tests: open field test, elevated plus maze test, modified three-chamber social test, marble burying task, and rotarod test. A large cohort of mice were used for all behavioral tests in the battery, except that a few mice died suddenly before the completion of the whole battery of tests. In addition, a second cohort of Sst-Pten mice were used for the social test only to balance the side preference, and a third cohort of mice were used for the home-cage activity test. Animals’ activity during the behavioral assessments was recorded by a Logitech C920 HD Pro Webcam, with the Ethovision software (Noldus, RRID: SCR_002798) used for further offline analysis. All experiments were conducted during the light cycle with room lights at around 80 lux except for the elevated plus maze (around 8 lux). All areas were cleaned with 70% ethanol between test sessions, including areas untouched by mice.

### Open field test

Mice were placed in the center of an open field apparatus (40 × 40 cm) and allowed to explore the field for 30 min. The movement was tracked using the Ethovision XT software. Distance moved and time spent in the center area (20 × 20 cm, square) and corner area (15 × 15 cm, triangle) were further analyzed. Self-grooming behaviors were also analyzed from the open field activity using mouse behavior recognition (Detection settings for each mouse: posture between 70-90%, probability higher than 75%) in the Ethovision XT.

### Home-cage activity test

Home-cage activity was recorded and analyzed using Ethovision XT software. Mice were individually housed in a clean cage for at least 24 h for acclimation. Mouse behaviors were recorded in its home cage with the filter top removed for 30 min. The distance moved was determined by off-line analyses of the video recordings.

### Elevated plus maze test

The elevated plus maze was composed of two open arms (30 × 5 cm) and two closed arms (30 × 5 cm) with 15 cm wall height and 50 cm above the floor. Mice were individually placed in the center of the maze and facing one of the open arms. They were allowed to explore the open and closed arms of the maze for 10 min. The duration of time in arms was recorded. The percentage of time in open arms (or closed arms) was calculated as the duration in open arms (or closed arms) / total exploring duration.

### Modified three-chamber social test

We used a modified apparatus to examine the social behaviors [79]. The apparatus [40 (H) × 60 (L) × 20 (W) cm] was divided into three chambers (20 cm long for each chamber) made from white Plexiglass. The side chambers and the center chamber were separated by white Plexiglass walls with a small gate (10 cm) in the middle. Square interaction areas (10 × 10 cm) in each side chamber was formed by the gate and a three-sided fence made from clear Plexiglass (Figure 1A). Test mice entered these two areas to interact with social partners. The two side chambers were covered by a transparent Plexiglass sheet during the three-session test. All three sessions were performed consecutively on one day. In the first session (habituation), the test mouse was placed in the center chamber and allowed to explore the chamber for 10 min. At the end of the habituation session, the test mouse was gently guided back to the middle of the center chamber, and the small gates were blocked. During the second session, which was to test the sociability, three novel C57BL/6 mice and three objects were placed in one of the two side chambers, respectively. The side chamber for social partners or objects was randomly assigned by flipping a coin. The test mouse was allowed to explore the center chamber for 10 min. After that, the test mouse was guided back to the middle of the center chamber again. In the third session, which was to test the social novelty preference, the objects were replaced by another set of three novel C57BL/6 mice, and the test mouse was allowed to explore the chamber for another 10 min. The duration that the animal spent in the interaction areas during each session was scored using the EthoVision XT software (Noldus). The preference index was calculated as the duration exploring the novel mice / the total duration exploring the objects (or familiar mice) and the novel mice.

### Marble burying task

Twelve glass marbles (3 × 4 arrangement) were placed on the surface of the bedding (5 cm deep) in standard mouse cages. The cages were covered with filter top during the test. Each test mouse was placed in the cage and allowed to explore for 30 min. The number of buried marbles was counted manually. The marble was checked from side of the cage. If the marble was buried more than 2/3 deep, it was counted as buried.

### Rotarod test

The two-session rotarod test was conducted on an accelerating rotarod (Panlab, Harvard Apparatus) over two days. Mice were allowed to become acclimated to the stationary rod for 60s on the first day. Each test session consisted of three trials with 15-min inter-trial intervals (ITI). During each trial, mice were placed on the rotating rod (4 rpm) facing away from the direction of rotation. The rotation speed was accelerated from 4 to 40 rpm over 5 min. If the mouse fell off the rod within 10 sec, the trial was repeated after 15 min. The latency to fall was recorded as the time delay between the start of the trial and the moment when mice fell off the rod or made a complete revolution on the rod.

### Statistical Analysis

Data was analyzed with SPSS (v.20) and GraphPad Prism 9 Software. All datasets were tested for normality using the Shapiro-Wilk test. For datasets with normal distribution, two-tailed *t*-test was used for comparisons between two groups, and two-way ANOVA with repeated measures in one factor (rmANOVA) or one-way ANOVA was used for comparisons among three groups. Tukey’s post hoc test was used for multiple comparisons after ANOVA test. For datasets with non-normal distribution, Wilcoxon matched-pairs signed rank test (Wilcoxon test) was used for comparisons between two groups (paired), and Kruskal-Wallis test with Dunn’s post hoc test was used for comparisons among three groups. One sample *t*-test or Wilcoxon signed-rank test was used for comparing the mean (or median) of a dataset with a hypothetical number. *p*-value < 0.05 was considered to be statistically significant.

## LIST OF ABBREVIATIONS

ASD: Autism spectrum disorder
PV: Parvalbumin
Sst: somatostatin
PTEN: Phosphatase and tensin homolog on chromosome ten
PI3K/AKT/mTOR: Phosphatidylinositol 3-kinase/AKT/mammalian target of rapamycin
KO: Knockout
WT: Wildtype
PFC: Prefrontal cortex
mPFC: Medial prefrontal cortex
vHipp: Ventral hippocampus
PC: Purkinje cells

## DECLARATIONS

### Ethics approval and consent to participate

Not applicable.

### Consent for publication

Not applicable.

### Availability of data and materials

The datasets used and/or analyzed during the current study are available from the corresponding author on reasonable request.

### Authors’ contributions

SS and AS collected the behavioral data. SS, AS and SH performed the data analyses. AS conducted the immunohistochemistry and western blot. SS, AS and SH wrote the manuscript. SH conceived and designed the experiments with the help of SS and AS. All authors read and approved the final manuscript.

## Acknowledgements

We thank Dr. John Hussman, Dr. Gene Blatt, and Ms. Elizabeth Benevides for critical reading of the manuscript. We also thank Drs. Louis DeTolla and Turhan Coksaygan at the University of Maryland School of Medicine for providing veterinary and consulting services.

---

**Figure S1.**
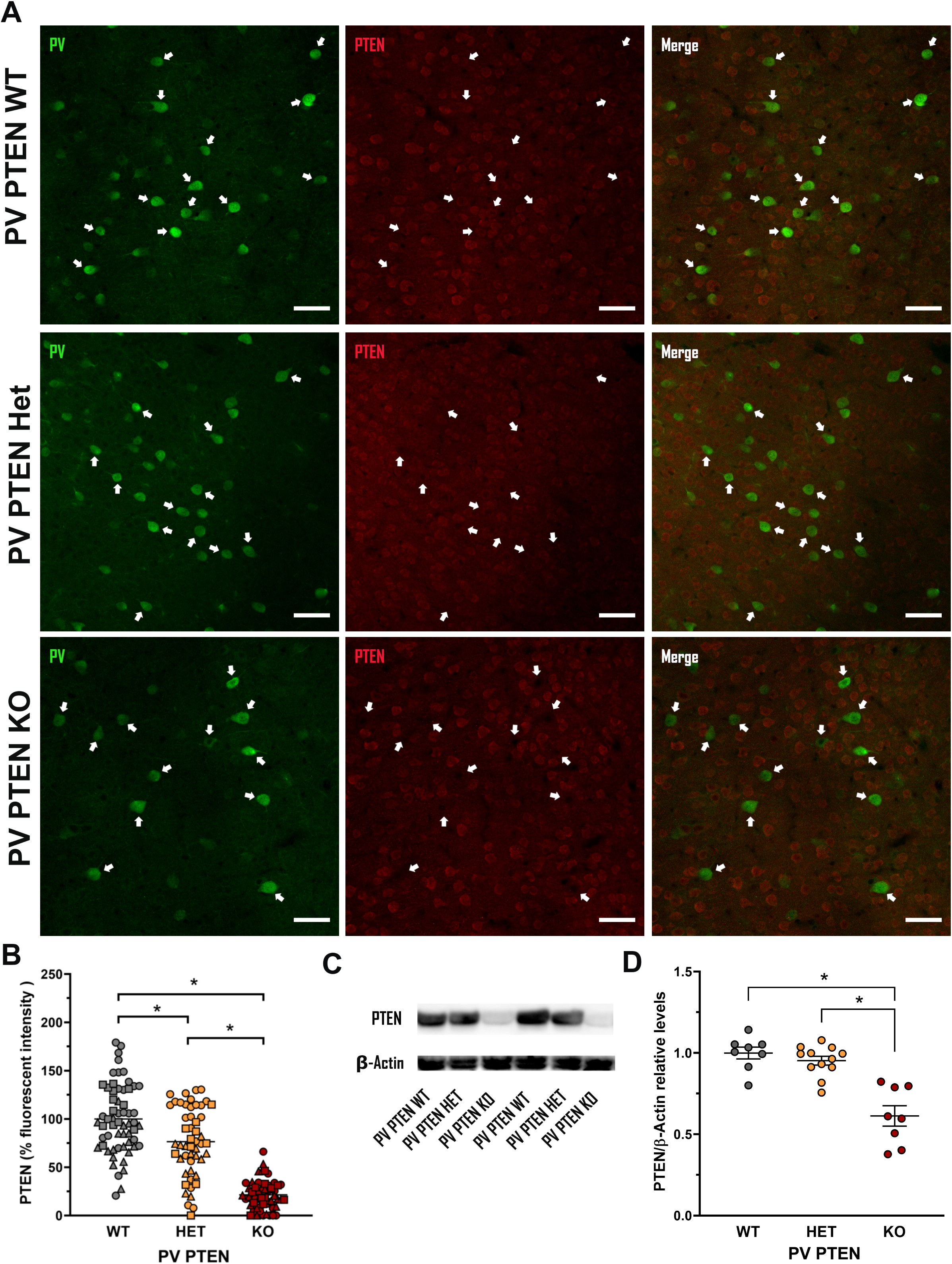
PTEN expression in parvalbumin positive neurons of PV-Pten mice. (**A**) Complete (PV-Pten-KO) or partial (PV-Pten-Het) genetic deletion of *Pten* in PV cells results in the absence or reduction, respectively, of PTEN immunoreactivity in PV^+^ cells, as shown in these cortical sections. Arrows point to PV^+^ cells. Scale bars represented 50 μm. (**B**) Quantification of PTEN fluorescence intensities in PV^+^ neurons normalized to WT mice (WT: n = 60 cells / 3 mice, Het: n = 51 cells / 3 mice, KO: n = 58 cells / 3 mice; Kruskal-Wallis statistic = 97.77, *p* < 0.0001, Kruskal-Wallis test; WT vs. Het: *p* = 0.0413, WT vs. KO: *p* < 0.0001, Het vs. KO: *p* < 0.0001, Dunn’s post hoc test). Each symbol shape represents a different animal. (**C-D**) Western Blot analysis showed decreased PTEN expression in PV-Pten-KO mice (WT: n = 8 mice, Het: n = 12 mice, KO: n = 8 mice; *F*_(2_, _25)_=23.90, *p* < 0.0001, one-way ANOVA; WT vs. KO: *p* < 0.0001, Het vs. KO: *p* < 0.0001, Tukey’s post hoc test). *: significant.

**Figure S2.**
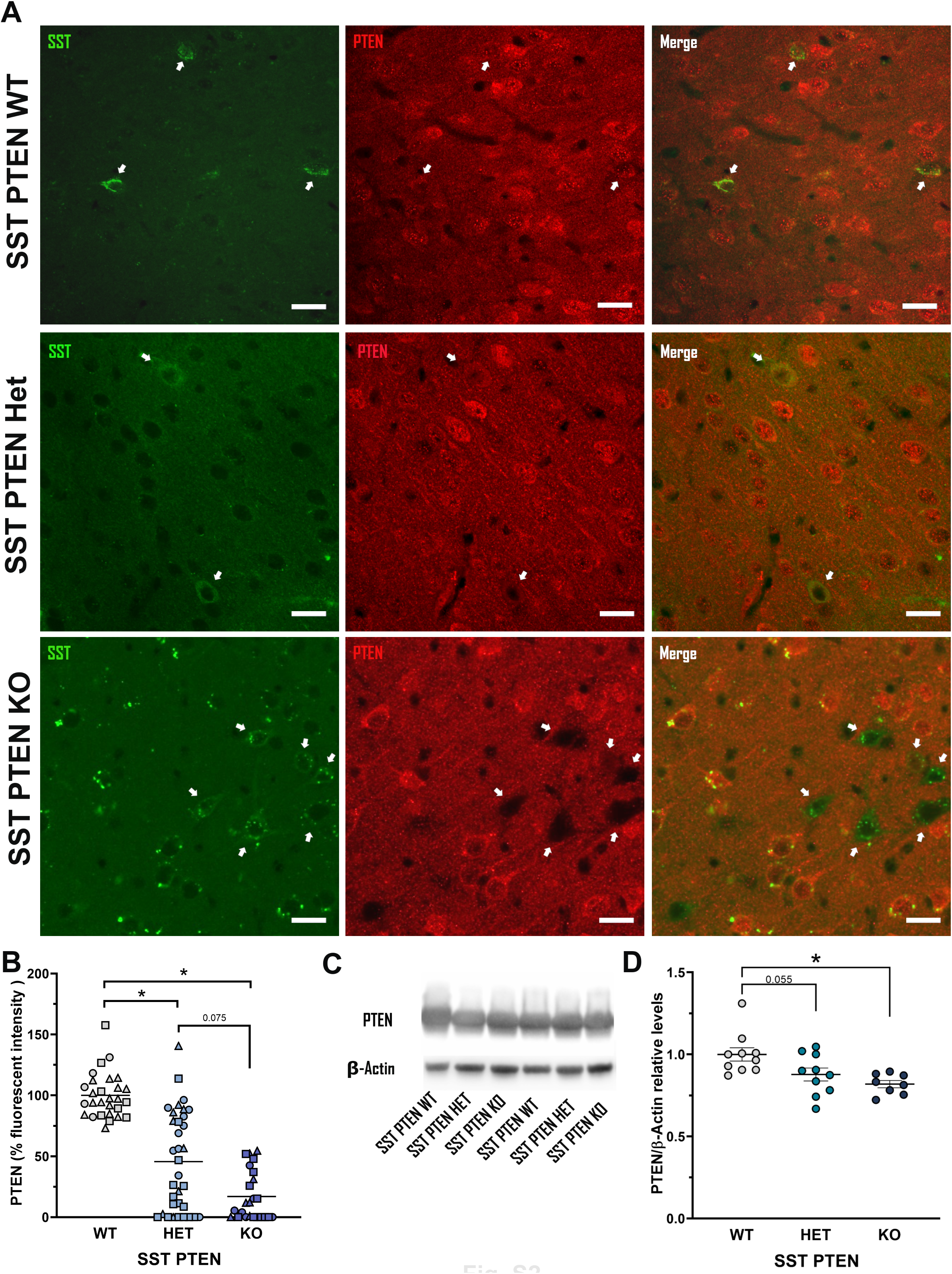
PTEN expression in somatostatin positive neurons of Sst-Pten mice. (**A**) Complete (Sst-Pten-KO) or partial (Sst-Pten-Het) genetic deletion of *Pten* in Sst cells results in a significant decrease of PTEN immunoreactivity in cortical Sst^+^ cells. Arrows point to Sst^+^ cells. Scale bars represented 25 μm. (**B**) Quantification of PTEN fluorescence intensity in Sst^+^ neurons as a percentage of WT PTEN ^+^ cells (WT: n = 28 cells / 3 mice, Het: n = 34 cells / 3 mice, KO: n = 24 cells / 3 mice; Kruskal-Wallis statistic = 46.48, *p* < 0.0001, Kruskal-Wallis test; WT vs. Het: *p* < 0.0001, WT vs. KO: *p* < 0.0001, Het vs. KO: *p* < 0.0752, Dunn’s post hoc test). Each symbol shape represents a different animal. (**C-D**) Western Blot analysis showed decreased PTEN expression in Sst-Pten-KO mice (WT: n = 10 mice, Het: n = 10 mice, KO: n = 8 mice; *F*_(2_, _25)_ = 6.26, *p* = 0.0063, one-way ANOVA; WT vs. Het: *p* = 0.0550, WT vs. KO: *p* = 0.0061, Tukey’s post hoc test). *: significant.

**Figure S3.**
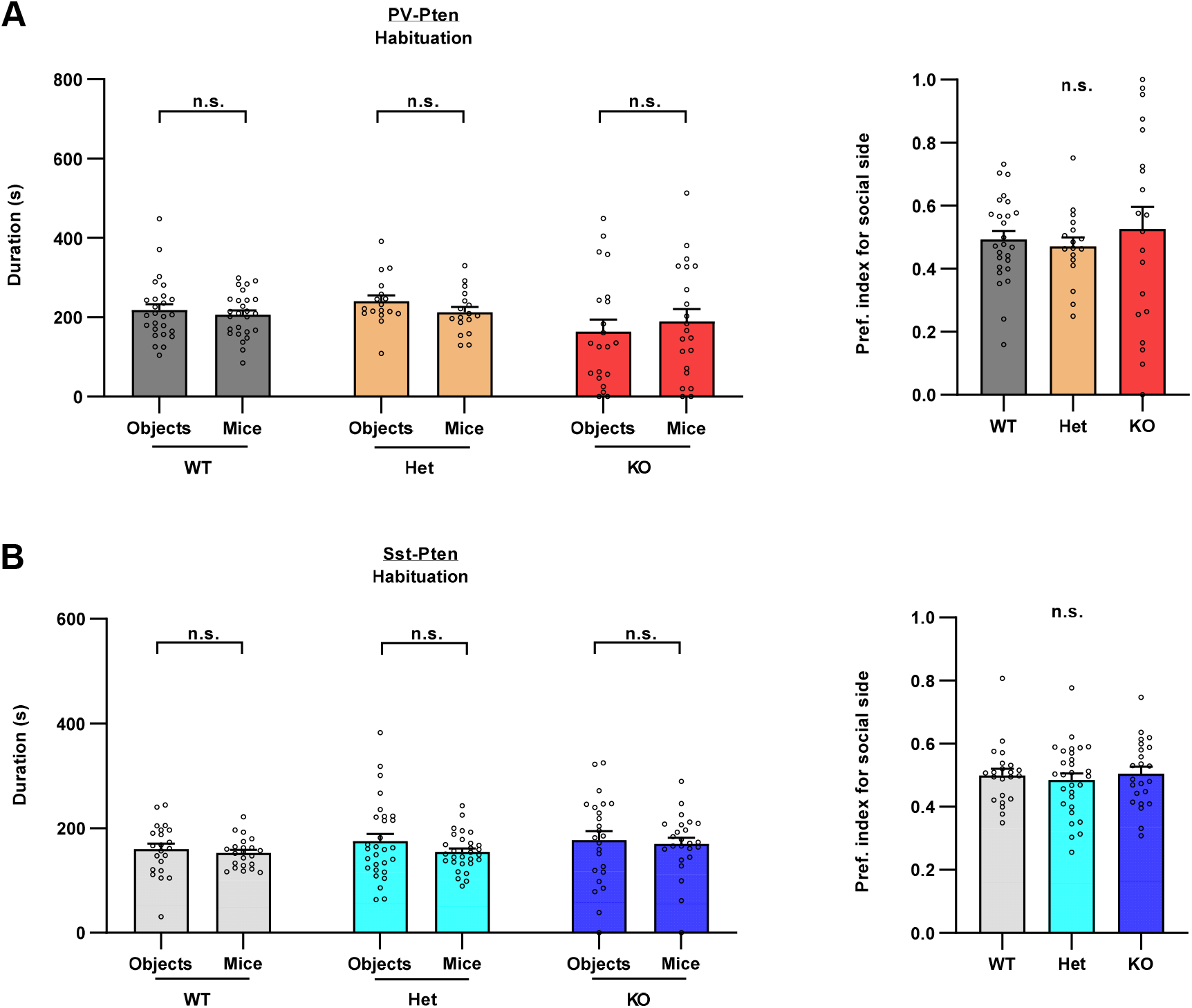
No pre-existing side preference during the habituation session in modified three-chamber social tests for both PV-Pten and Sst-Pten mice. **(A)** PV-Pten mice spent similar time in the interaction zones that would be assigned to social partners (Mice) or objects (WT: n = 26, *p* = 0.8809, Wilcoxon test; Het: n = 17, *p* = 0.3168, paired *t*-test; KO: n = 21, *p* = 0.6320, paired *t*-test). Their preference indexes for social side are not statistically different from 0.5 (WT: *p* = 0.7837, Het: *p* = 0.3217, KO: *p* = 0.7170, one-sample t test). **(B)** Sst-Pten mice spent similar time in the interaction zones that would be assigned to social partners (Mice) or objects (WT: n = 22, *p* = 0.5159, paired *t*-test; Het: n = 29, *p* = 0.2070, paired *t*-test; KO: n = 24, *p* = 0.6650, paired *t* test). Their preference indexes for social side are not statistically different from 0.5 (WT: *p* = 0.8987, Wilcoxon signed rank test; Het: *p* = 0.4937, one sample *t* test; KO: *p* = 0.8282, one-sample *t* test). The preference index for social side was calculated as the duration exploring the social side / the total duration exploring the social side and the object side. n.s.: not significant.

**Figure S4.**
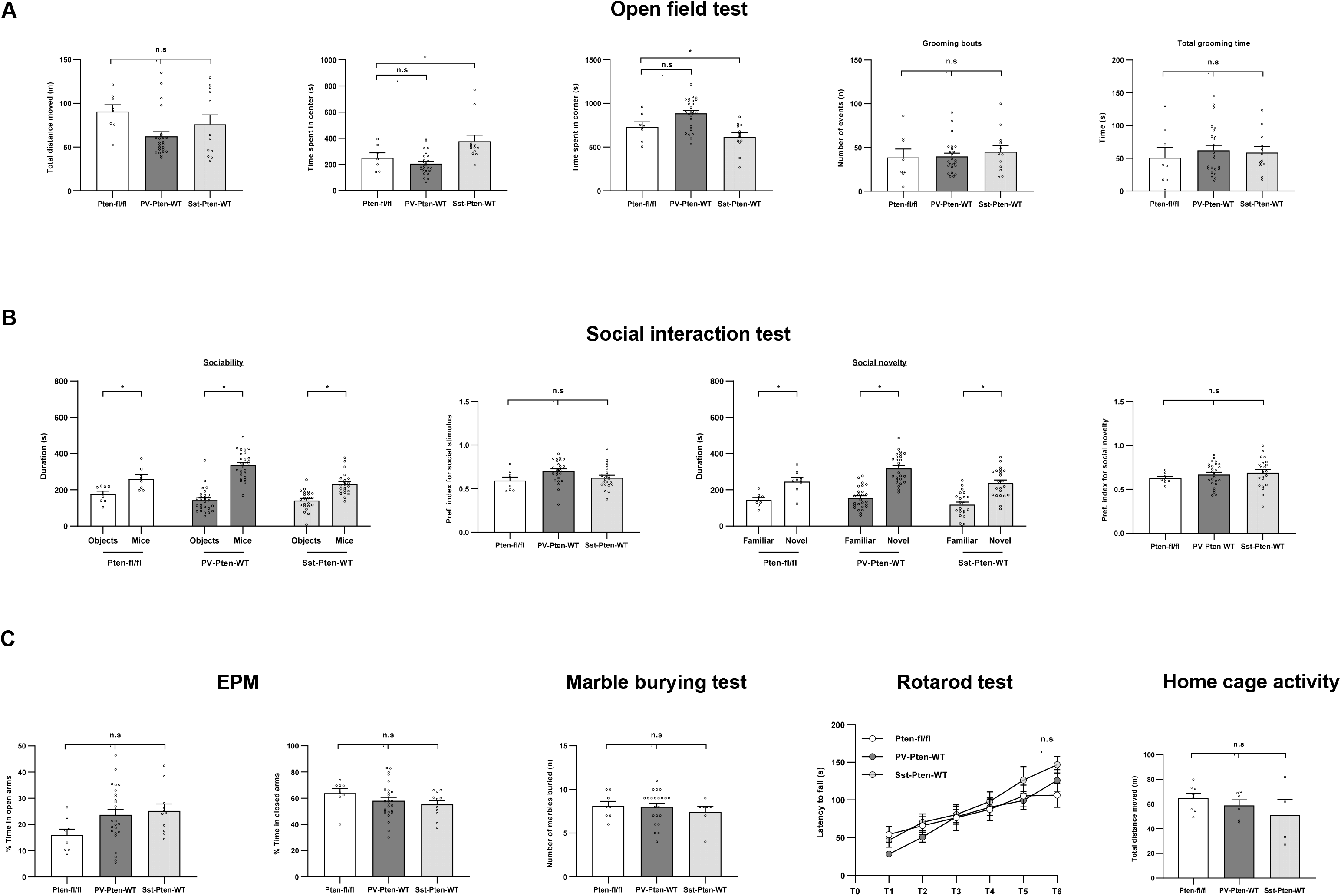
Cre expression in PV- or Sst-neurons did not change most behavioral performance. (**A**) No significant change in open filed test was observed in PV-Pten-WT or Sst-Pten-WT mice compared with Pten^fl/fl^ mice, except that Sst-Pten-WT displayed increased time spent in the center of the open field. (**B**) No significant change in social interaction test was observed in PV-Pten-WT or Sst-Pten-WT mice. (**C**) No significant change in EPM, marble burying test, rotarod test or home-cage activity was observed in PV-Pten-WT or Sst-Pten-WT mice. Data are represented as mean ± SEM. *: significant; n.s.: not significant.

**Figure S5.**
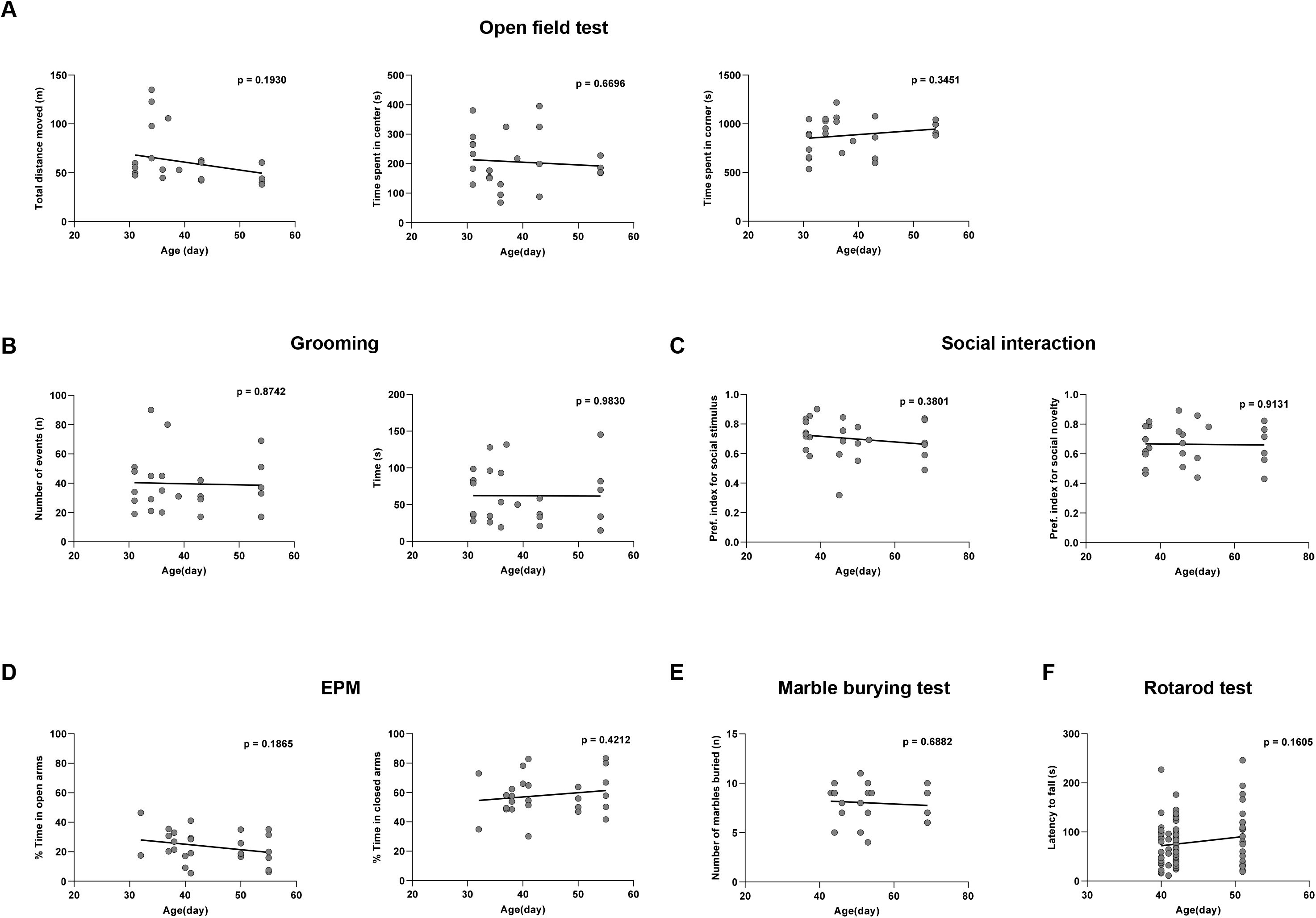
No significant correlation between behavioral performance and ages in PV-Pten-WT mice. Pearson correlation tests were performed between ages and the behavioral output in open fieled test (**A**), self-grooming (**B**), social interaction (**C**), elevated-plus maze test (EPM) (**D**), marble burying test (**E**), and rotarod test (**F**) in PV-Pten-WT mice. None of the correlation was significant.

**Figure S6.**
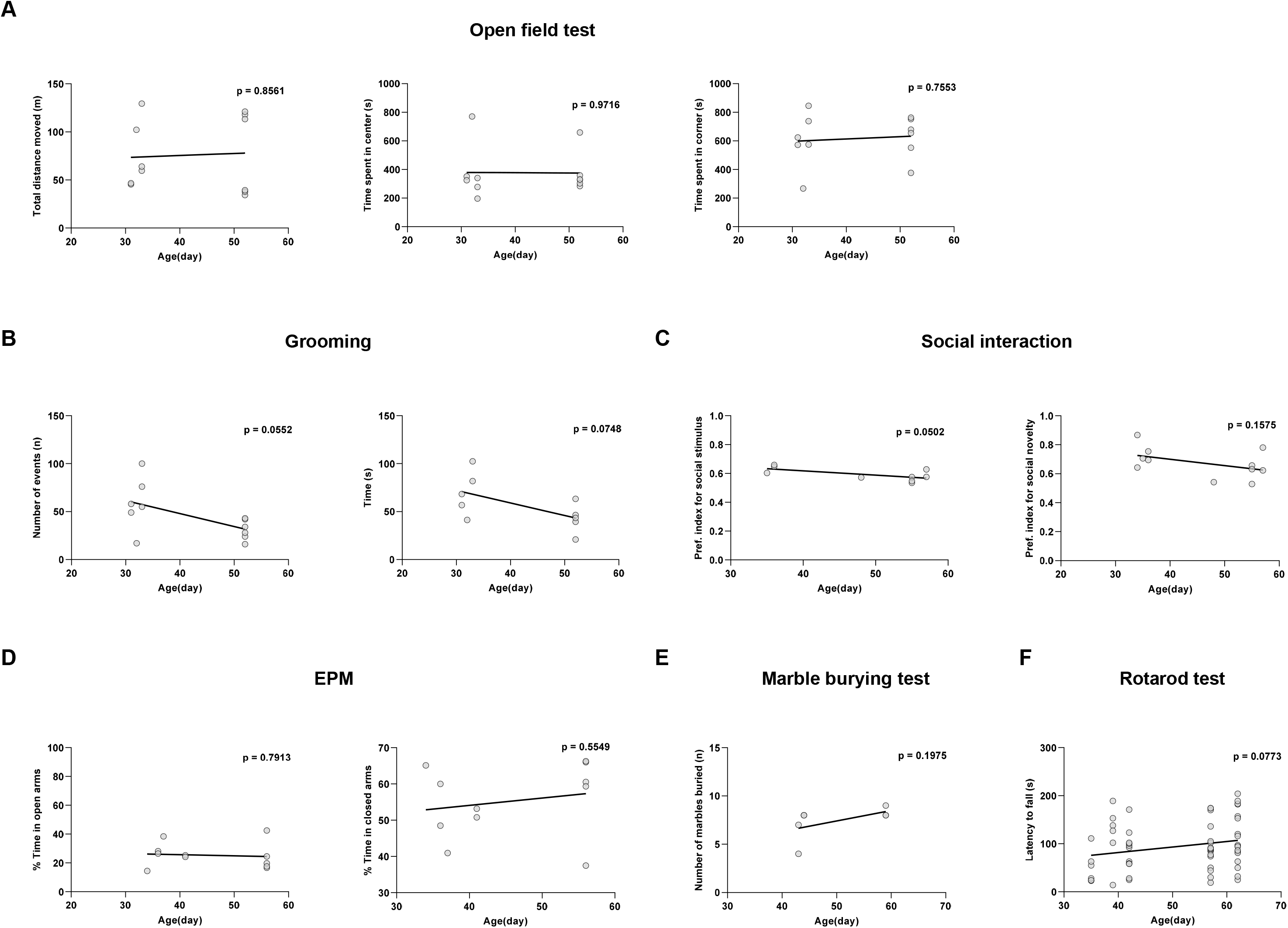
No significant correlation between behavioral performance and ages in Sst-Pten-WT mice. Pearson correlation tests were performed between ages and the behavioral output in open field test (**A**), self-grooming (**B**), social interaction (**C**), elevated-plus maze test (EPM) (**D**), marble burying test (**E**), and rotarod test (**F**) in Sst-Pten-WT mice. None of the correlation was significant.

**Figure S7.**
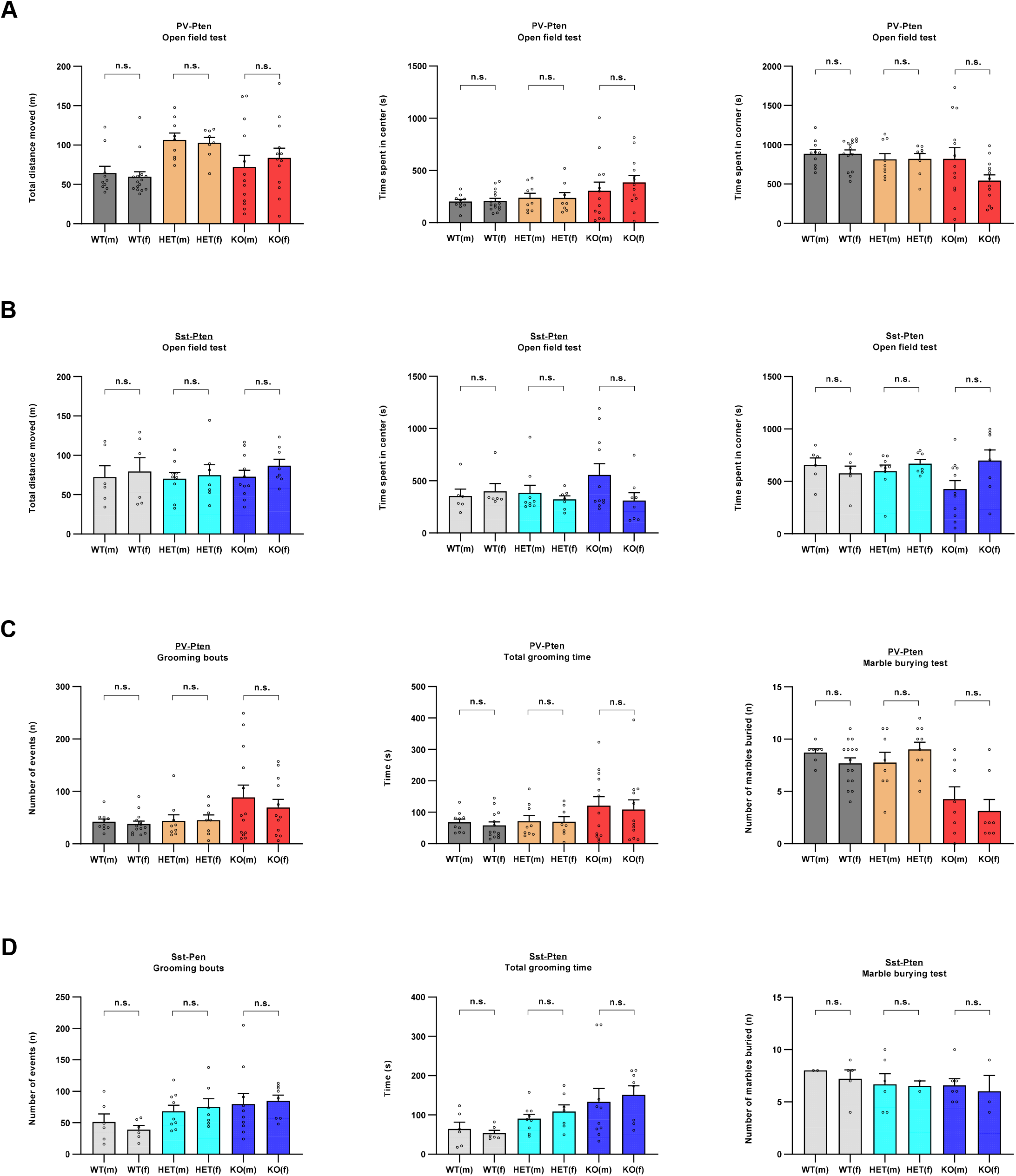
No sex effect in open-field test and marble burying test was observed in PV-Pten or Sst-Pten mice. **(A)** There was no sex effect in open-field test in PV-Pten mice. **(B)** There was no sex effect in open-field test in Sst-Pten mice. **(C)** There was no sex effect in grooming and marble burying test in PV-Pten mice. **(D)** There was no sex effect in grooming and marble burying test in Sst-Pten mice. Data are represented as mean ± SEM. n.s.: not significant.

**Figure S8.**
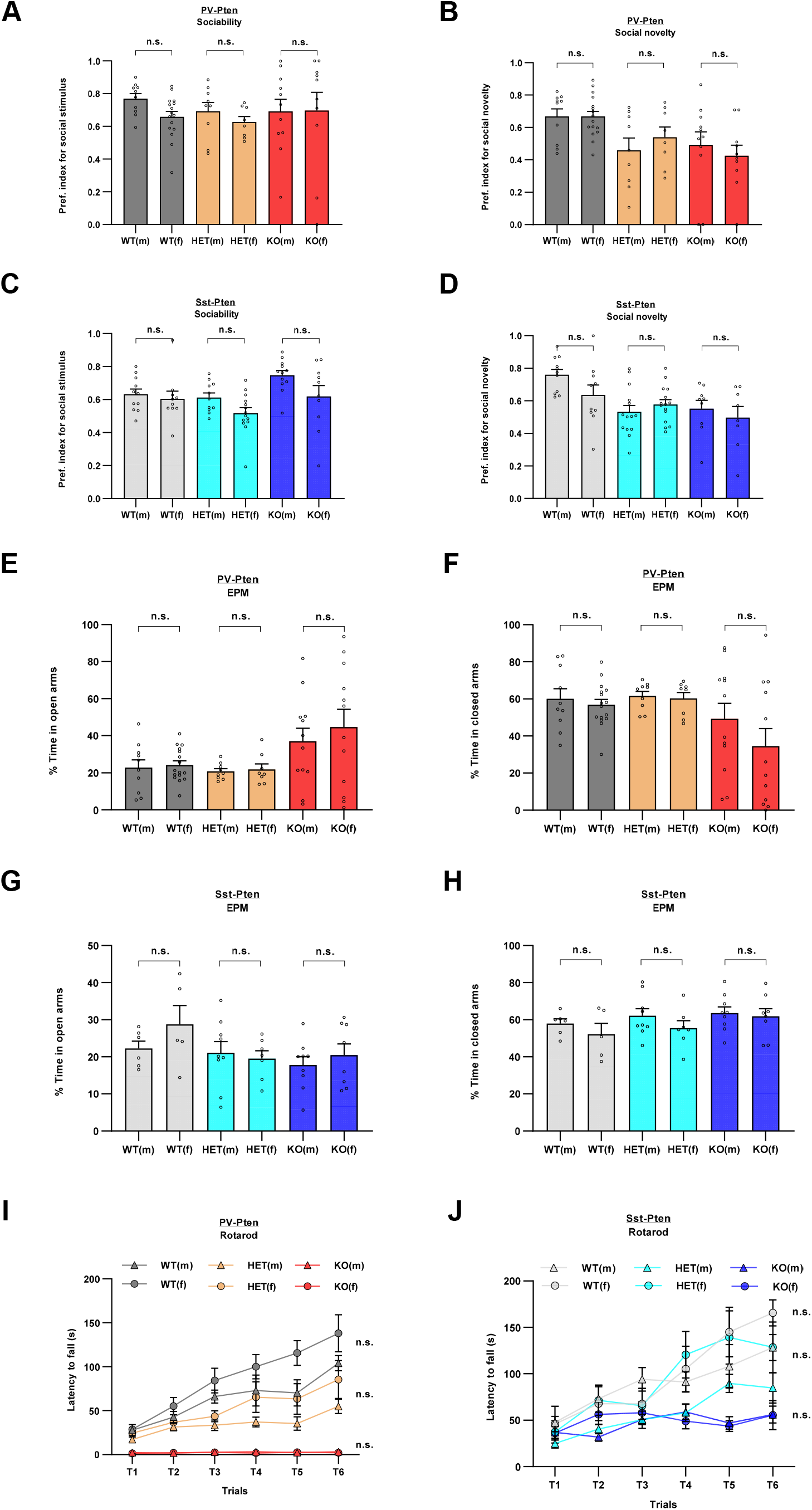
No sex effect in social test, rotarod test and EPM test was observed in PV-Pten or Sst-Pten mice. (**A-B**) There was no sex effect in sociability (A) or social novelty preference (B) in PV-Pten mice. (**C-D**) There was no sex effect in sociability (C) or social novelty preference (D) in Sst-Pten mice. (**E-F**) There was no sex effect in elevated-plus maze (EPM) test in PV-Pten mice. (**G-H**) There was no sex effect in elevated-plus maze (EPM) test in Sst-Pten mice. (**I**) There was no sex effect in rotarod test in PV-Pten mice. (**J**) There was no sex effect in rotarod test in Sst-Pten mice. Data are represented as mean ± SEM. n.s.: not significant.

